# Downscaling global reference points to assess the sustainability of local fisheries

**DOI:** 10.1101/2023.12.15.571896

**Authors:** Jessica Zamborain-Mason, Sean R. Connolly, M. Aaron MacNeil, Michele L. Barnes, Andrew G. Bauman, David A. Feary, Victor Huertas, Fraser A. Januchowski-Hartley, Jacqueline D. Lau, Michalis Mihalitsis, Joshua E. Cinner

**Affiliations:** College of Science and Engineering, James Cook University,Townsville, Queensland, Australia; Harvard T.H. Chan School of Public Health, Boston, MA 02115, USA; Smithsonian Tropical Research Institute, Panama City, Panama; Ocean Frontier Institute, Department of Biology, Dalhousie University, Halifax, NS B3H 3J5, Canada; College of Arts, Society and Education, James Cook University, Townsville, Queensland, Australia; Nova Southeastern University_Fort Lauderdale, FL, USA; MRAG Ltd, London, W1J 5PN, United Kingdom; Research Hub for Coral Reef Ecosystem Functions, James Cook University, Townsville, QLD, Australia; Department of Evolution and Ecology, University of California, Davis, CA, 95616

**Keywords:** MMSY, coral reefs, small-scale fisheries, stock assessment, scale-mismatch, fishers’ perceptions, Papua New Guinea

## Abstract

Multispecies coral reef fisheries are typically managed by local communities who often lack research and monitoring capacity, which prevents estimation of well-defined sustainable reference points to perform locally relevant fishery assessments. Recent global advances in modelling coral reef fisheries have developed pathways to use environmental indicators to estimate multispecies sustainable reference points. These global reference points are a promising tool for assessing data-poor reef fisheries but need to be downscaled to be relevant to resource practitioners. Here, using a small-scale multispecies reef fishery from Papua New Guinea, we estimate sustainable reference points and assess the sustainability of the fishery by integrating global-scale analyses with local-scale environmental conditions, fish catch, reef area, standing biomass estimates, and fishers’ perceptions. We found that assessment results from global models applied to the local context of our study location provided results consistent with local fishers’ perceptions. Specifically, our downscaled results suggest that the fishing community is overfishing their reef fish stocks (i.e., catching more than can be sustained) and stocks are below B_MMSY_ (i.e., below biomass levels that maximize production), making the overall reef fishery unsustainable. These results were consistent with fisher perceptions that reef fish stocks were declining in abundance and mean fish length, and that they had to spend more time finding fish. Our downscaled site-level assessment reveals severe local resource exploitation, whose dynamics are masked in national-scale assessments, emphasizing the importance of matching assessments to the scale of management. More broadly, our study shows how global reference points can be applied locally when long-term data are not available, providing baseline assessments for sustainably managing previously un-assessed multispecies reef fisheries around the globe.

## Introduction

Multispecies coral reef fisheries are a major source of food and nutrients for many tropical coastal communities around the globe. Yet, the sustainability of many of these fisheries remains unassessed at scales relevant for fisheries management, mainly due to a lack of clearly defined, locally relevant sustainable reference points against which fishery performance can be assessed (Branch et al. 2011). Estimating sustainable reference points—such as multispecies maximum sustainable yield (MMSY) or the standing biomass at which MMSY is achieved (B_MMSY_)—for a given location usually requires reliable fisheries statistics obtained from long-term monitoring (Worm et al. 2009) and information on how fish populations naturally recover (McClanahan 2018). However, multispecies coral reef fisheries are typically data-poor and predominantly managed locally (e.g., local communities), where monitoring and management capacity may be lacking (Mora et al. 2009; Worm and Branch 2012; Darling and D’Agata 2017; Samoilys et al. 2017). Thus, for such a fishery, using only locally available data can severely limit the extent to which relevant sustainable reference points can be estimated.

Recent findings from global analyses of coral reef fisheries can provide additional information that, leveraged along with local data, may facilitate (i) the estimation of sustainability benchmarks and (ii) performing assessments at local management scales. The compilation of large global datasets of reef fish biomass (i.e., fisheries-independent surveys, e.g., McClanahan et al. 2011; Graham et al., 2017; Cinner et al. 2020) has enabled estimation of sustainable reference points for multispecies coral reef fisheries using local environmental conditions (e.g., Zamborain-Mason et al 2023). These, together with reconstructed fish landings (Zeller et al. 2016) and reef area estimates (UNEP-WCMC 2010), have allowed the assessment of multispecies coral reef fisheries over relatively large geographical scales (e.g., country assessments). Yet large-scale assessments alone are inadequate to effectively inform resource practitioners at the scales of reef fisheries management (e.g., Cash and Moser 2000). Uncertainty about fishery status for a given nation (e.g., Zamborain-Mason et al. 2023), spatial heterogeneity in reference points, catch, and standing biomass within national borders (e.g., Zeller et al. 2016; Karisa et al. 2020), as well as local exploitation context all impact local sustainability (Kerr et al. 2017; McClanahan and Kosgei 2023). Taking advantage of global information requires the scale of assessment to match with the scale of management (Cash and Moser 2000; Hibbard and Janetos 2013; McClanahan and Kosgei 2023).

Here, by integrating global model outputs (Zamborain-Mason et al. 2023) with local scale environmental and fisheries information, we demonstrate how recent global reference points for coral reef fish stocks can be downscaled to inform local fisheries management when local fisheries-relevant data are insufficient by themselves for a sustainability assessment (e.g., no fish population recovery data). Using a previously unassessed small-scale multispecies coral reef fishery from Papua New Guinea, we (i) estimate local MMSY sustainable reference points for the fishery; (ii) assess the status of local reef fish stocks relative to these key reference points; (iii) quantify how much the long-term food-provisioning potential of local reefs could increase if stocks were managed sustainably, and how long it would take to reach such a state; and (iv) compare assessment results with local fishers’ perceptions. Together, our study shows how global reef fisheries benchmarks can be applied locally, thus providing a tool to assess and sustainably manage un-assessed reef fisheries around the globe at a scale relevant for management.

## Methods

### Study site and fishery context

We conducted fieldwork in Ahus Island, a coastal island of approximately 950 people, who are highly dependent on reef resources (Barnes et al. 2022), in Manus Province, Papua New Guinea (Fig. 1). Historically, islanders managed their reefs through customary systems (Cinner 2005): clan leaders and individuals with sea tenure rights applied temporary fishing closures, and had rights to certain fishing practices (e.g., gears) and specific operating times (i.e., night fishing). However, in recent years, the customary system has eroded and compliance with customary restrictions has faded as population has increased (Lau et al., 2020). The reef fishery lacks MMSY sustainable reference points, has not been assessed against locally-specific benchmarks and has no current fisheries management measures in place.

**Figure 1.**
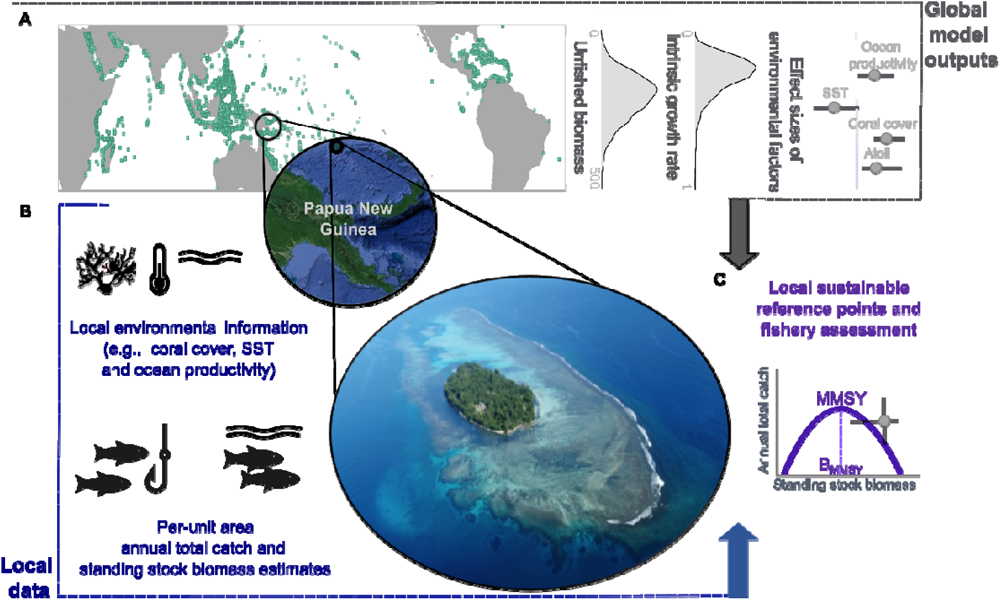
Downscaling global benchmarks to local fisheries: Study setting. **(A)** Global map of tropical coral reefs (UNEP-WCMC 2010) showing our study site and reef area. Distributions are examples of the posteriors from global model outputs (Zamborain-Mason et al. 2023) that are used to apply to local context: unfished biomass of reef fish (115.6 [97.5 - 140.6] t/km^2^; (posterior median [90% posterior uncertainty intervals]); community growth rate (0.14 [0.08 - 0.31] year^-1^) and effect sizes (i.e., slopes in a linear model) of environmental covariates on log unfished biomass (coral cover: 0.69 [0.41 - 0.97] ; ocean productivity: 0.43 [0.09 – 0.75]; SST: -0.35 [-0.74 – 0.08]; and Atoll: 0.49 [0.14 – 0.81])**. (B)** Type of local-scale information collected to downscale global estimates: Local environmental information to estimate sustainable reference points, and per-unit-area standing stock biomass and annual catch estimates to assess the status of stocks. **(C)** Example of outputs from downscaling global models to local context to perform fishery assessments (e.g., local surplus production and reference points (purple) combined with estimates of standing biomass and annual total catch (gray)). Photo credit: Dean Miller.

As with many tropical small-scale reef fisheries (e.g., Grantham et al. 2021), fishing patterns (e.g., catch volumes, composition, and locations) in Ahus are driven by seasonal weather cycles, namely the windy and the non-windy season. Ahus’s reef fisheries are opportunistic and multispecies. Fishers use a range of gears such as spear guns, trolling lines, hand lines, nets, and hand spears. Some households own or rent motorized vessels to target pelagic fishes such as tuna; however most male fishers use hand-paddled canoes and target reef-associated fishes. Motorized vessels are sometimes used to fish reef-associated species, for instance for cultural ceremonies. Gleaning is predominantly undertaken by women and children, focusing on lagoon and backreef area habitats, with invertebrates and reef fishes targeted.

### Data collection and analyses

To estimate sustainable reference points and assess the sustainability status of the multispecies reef fishery using global model outputs we combined three types of local information: (i) environmental characteristics (i.e., coral cover, ocean productivity, sea surface temperature (SST), and whether the location is an atoll); (ii) per-unit area estimates of standing reef fish biomass; and (iii) per-unit-area estimates of annual reef fish catch (Fig. 1).

#### Environmental context

We sampled a total of twelve reef sites spanning depths from three to ten meters, in slope and lagoon reef habitats, across four different years (2009, 2012, 2016, 2018). Overall, estimates of coral cover, sea surface temperature (SST) and ocean productivity for our study were obtained by averaging site and year-specific measurements or estimates. Our study location was characterized as non-atoll. Live hard coral cover at each site was recorded using replicate 4 × 50 m (2009) or 6 × 30 m (2012, 2016 and 2018) point intercept methods. Transects were laid parallel to the reef crest and the substrate directly beneath the transect tape was surveyed every 0.5 m (2009, 2012 and 2016) or 1 m (2018) across all transects. Remotely sensed monthly net primary production (i.e., ocean productivity; Behrenfeld and Falkowski 1997;) and SST estimates (Boyin et al. 2017) for each of our sites were annually averaged.

#### Per-unit-area standing reef fish biomass

Reef fish biomass estimates were recorded through underwater visual census (UVC) at the same sites as coral cover using belt-transects. Consistent with the data used to develop global models (i.e., Zamborain-Mason et al. 2023), diurnally-active, non-cryptic reef fishes above 10 cm length from families resident on the reef (Table S1) were counted, identified to species level, and had their total length estimated. Total observed biomass of fishes on each transect was calculated using published species-specific length-length and length–weight relationships (http://fishbase.org; Froese and Pauly 2018; Boettiger et al. 2012). When parameters were not available for specific species, we used parameters for a closely related and similar sized species. Site-specific per-unit area biomass for each year was estimated by summing the observed biomass from all transects within a site and dividing by the number of transects performed. To make site-specific biomass data comparable to the global models, observed site biomass data were corrected for sampling and methodological effects using the median posterior effect sizes from the global models (i.e., standardizing our site’s depth, sampled area and habitat to global average conditions (i.e., 6.7m and 518 m^2^ and slopes), and subtracting/adding the relevant effects to observed biomass values; Zamborain-Mason et al. 2023; Fig. 1). As the sampled sites varied among years and seasons, we used the entire distribution of standing stock biomass estimates from all sites and years (n=26) and estimated the median standing stock and associated 95% adjusted confidence intervals using non-parametric bootstrapping with replacement for 4000 iterations (Davison and Hinkley 1997; Canty 2002).

#### Per-unit-area annual catch

Catch per unit area for our study location was obtained by dividing annual reef catch estimates by reef area. Total reef area used by local fishers (∼9 km^2^) was estimated using the Millennium Mapping project dataset (IMaRS-USF 2005). Annual reef fish catch estimates were obtained by combining (i) catch surveys collected in two points in time representing different seasons (May-June 2018 and February 2019), and (ii) household structured social surveys collected in May 2018.

Catch surveys (Table S2) were performed at landing sites (i.e., approaching fishers as they returned from fishing activities), individual households (e.g., if a fisher had recently returned from fishing) or at local markets. All catch surveys involved: (i) photographing the catch against a size scale (Cinner and McClanahan 2006), (ii) recording the gear used, boat type, the number of fishers, their biological sex, the effective time spent fishing or effort (i.e., fishing trip hours minus the travelling time where the gear was not deployed), the destination of the catch, and the fishing grounds (i.e., fishers were presented with a map of the study site and asked to identify where the catch came from), and (iii) questions about long-term effort (answered only once by each recorded individual fisher who had time to answer the long-term questions). We typically observed fishers while they were fishing, corroborating their reported fishing grounds. We also made sure fish were not double counted by obtaining fishing trip details at the start of all catch surveys (i.e., fisher, start and end time of fishing trip, and hours that gears were employed). Catch photographs were analysed to identify individual fish to the lowest taxonomic level possible and measure their standard length. Biomass was estimated as outlined above for the standing stock biomass, using published species-specific length-length and length–weight relationships (Froese and Pauly 2010). Note some of the photographs were of processed fish (e.g., smoked or fried). Cooking tends to decrease the length and weight of a fish. Although the effect on overall catch biomass is likely negligible, catch estimates from such photographs may have led to small downward biases in some cases.

A total of 428 fishing trips, 203 individual fishers and five different gears (handline, speargun, trolling line, simple spear, and gillnet) were recorded during our surveys. Catch composition included reef associated-fish, invertebrates, sea turtles, and other fish (e.g., tuna and sharks). As we were interested in reef assemblages and wanted to make estimates comparable to global reference points, for the catch and effort estimates we only used surveys with reef associated fish families (Table S1), where individual fish total length was above 10 cm and fish were caught in designated reef areas (n=340). Fisher-specific catch was estimated by dividing the total catch by the number of fishers on the fishing trip. For fishing trips that included a mix of target groups (e.g., pelagic and reef-associated fish) and where overall fishing grounds overlapped with the designated reef area, we estimated the proportion of reef-fish in the catch and assumed effort was proportional. This allowed us to estimate fisher-specific reef fish catch per unit effort (CPUE) distributions for each season. Median annual catch per fisher (and 95% adjusted confidence intervals) was estimated from fisher-specific CPUE and season-specific effort distributions using non-parametric bootstrapping with 4000 iterations (i.e., allowing for different subsample sizes). Keeping the data-structure (i.e., distributions of different sample sizes), for each sex category (i.e., female and male) we estimated (i) the median CPUE and median hours fishing for each season (i.e., windy and non-windy), and (ii) the median total annual catch per fisher, by dividing the year into the two six-month seasons. To examine potential bias in our fish catch sampling, we: (i) compared CPUE distributions from fishers that responded to the long-term questions and those that did not, and found they were similar (Fig. S1; Kolmogorov-Smirnov D statistic -0.09; p=0.85); and (ii) compared the CPUE distributions from fishers whose landings were observed more than once to fishers that were only recorded once, and also found they were similar (Fig. S1; D statistic -0.109; p=0.27). Note that informally, fishers reported that catch and effort during the survey period was typical for the season.

Household structured social surveys were used to extrapolate fisher-specific median total annual catch to total community annual catch estimates. We surveyed household heads (n = 138 out of 140 households) and recorded the livelihood activities of all members of the household, including fishing and gleaning, and the specific fishing gears used. This yielded a total of 152 male fishers and 131 female gleaners in the community. In order to estimate annual reef fish catch we excluded three male fishers who only reported using trolling line as their gear because trolling generally targets pelagic fish away from reef areas. Therefore, a total of 149 male and 131 female fishers targeting reef fishes were estimated for this study location. We multiplied sex-specific total annual catch times the number of fishers in each sex category to get annual catch estimates for our study location, and calculated catch per unit area by dividing the total annual catch by the estimated reef area.

#### Estimating local sustainable reference points from global models

Whole assemblage multispecies maximum sustainable reference points (i.e., MMSY and B_MMSY_) were obtained using local environmental conditions (i.e., coral cover, ocean productivity, SST, and the non-atoll nature of our study location) in combination with the posterior multispecies unfished biomass, the community biomass growth rate, and the effect sizes of environmental factors on unfished biomass from global models (Zamborain-Mason et al. 2023; Fig. 1). We used global estimates based on an aggregate Graham-Schaefer surplus growth model that treats the whole assemblage as a single stock (e.g., Mueter and Megrey 2006; Link 2017), and assumes a symmetric (logistic) population growth curve whose productivity (and thus sustainable yields) peaks at 0.5 of unfished biomass (Schaefer 1954). However, we also estimated sustainable reference points and assessed the fishery using other versions of the Pella-Tomlinson (Pella and Tomlinson 1969) surplus production models (of which the Graham-Schaefer is one special case): the Gompert-Fox for which production peaks below 0.5 of unfished biomass (Fox 1970; Tjorve and Tjorve 2017), and the Pella-Tomlinson with scale parameters of 3 and 4 for which production peaks at >0.5 of unfished biomass (Quinn and Derison 1999; Table S3). Additionally, we estimated reference points using site-specific environmental estimates for different years but found little variation (Fig. S2).

#### Local assessment of reef fish stocks

To provide a baseline assessment of the local multispecies reef fishery, per-unit-area standing biomass and total annual catch were compared to the estimated surplus production curve. Reefs were classified as (i) ‘below B_MMSY_’ if median standing biomass estimates were below the median B_MMSY,_ the biomass at which sustainable yields are expected to be maximized (Garcia et al. 2018), and (ii) as ‘undergoing overfishing’ if median total catch was above the median estimated surplus, given their standing biomass estimates (Hilborn 2011). The probability of being below B_MMSY_ and overfishing was also estimated incorporating uncertainty from the surplus production estimates. The fishery sustainability status was categorized following global assessments (Zamborain-Mason et al. 2023) as follows: in good condition (not below B_MMSY_ and not catching above MMSY), unsustainable (below B_MMSY_ and overfishing), warning (not below B_MMSY_ and catching above MMSY) or recovering (below B_MMSY_ and not overfishing). Additionally, we estimated the sustainable yield lost from assemblages below B_MMSY_ (i.e., potential gains from recovering stocks), as the difference between the whole assemblage MMSY and the estimated surplus. Minimum required biomass gains for recovery were estimated as the difference between the estimated B_MMSY_ and current median biomass levels. If uncertainty intervals returned negative values, we reported those values as zero (i.e., no biomass gains required). We repeated the above recovery assessment using a more conservative target, the biomass levels corresponding to ‘pretty good multispecies yields’ (i.e., B_PGMY_; which is where biomass levels correspond to the right-hand side of the surplus production curve that produces 0.8 of MMSY; Hilborn 2010; Rindorf et al. 2017 and is associated with increased diversity and ecosystem function; Zamborain-Mason et al. 2023). We then calculated location-specific recovery timeframes under a seascape moratorium scenario, corresponding to a scenario at which a fishing moratorium was imposed on the entire fishery (this estimates the minimum time for recovery to B_MMSY_ and B_PGMY_ for the location). These timeframes were calculated by combining the posterior community biomass growth rates from global models, with the locally estimated standing biomass, unfished biomass and B_MMSY_ or B_PGMY_ estimates. Assuming a logistic growth curve of whole assemblage biomass through time, the recovery time (Rt) is:

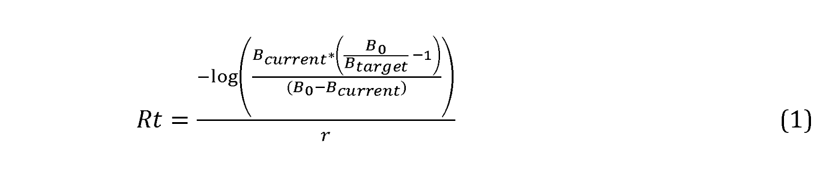

where B_current_ is the estimated standing biomass, r is the posterior community biomass growth rate from global models, B_0_ is the estimated posterior unfished biomass and B_target_ is the biomass benchmark or target to be achieved (i.e., B_MMSY_ or B_PGMY_). Similar to required biomass gains, if uncertainty intervals returned negative recovery times, we reported those values as zero.

#### Fishers’ perceptions

Finally, to assess how consistent our assessment results were with fishers’ perceptions, catch surveys also included a section about perceived reef fish stock status and drivers (Table S2), answered once by each recorded individual fisher willing to answer the long-term questions (n=77). Fishers were asked whether, compared to five years ago (i) the amount of reef-fish had increased, decreased or stayed the same, (ii) the size of reef-fish had increased, decreased or stayed the same, (iii) they had to spend more time traveling to catch reef fish, and (iv) they caught the same reef species, and if not, why.

## Results

Based on the Graham-Schaefer surplus production model, we estimate an 88% chance that reef fishers were catching above MMSY and a 61% chance that stocks are below B_MMSY_ (Fig. 2B), suggesting the multispecies reef fishery is unsustainable and reef fish populations are ongoing decline. Specifically, estimated multispecies maximum sustainable yields (MMSY) for the coral reef fishery at our study location were 1.9 [0.8 - 4.3] t/km^2^/y (median [90% posterior uncertainty intervals]), attained at standing biomass values, or B_MMSY_, of 18.3 [10.2 - 34.8] t/km^2^ (Fig. 2A). However, based on current biomass of 15.6 [12.6 – 22.4] t/km^2^ (median and 95% adjusted bootstrap confidence intervals), reefs are below B_MMSY_ (B<BMMSY), so the catch that can be sustained in the long-term is 1.6 [0.5 – 4.3] t/km^2^/y (e.g., ∼ 88% of MMSY). Estimated annual catch was 4.6 [1.6 – 8.7] t/km^2^/y (median [95% adjusted bootstrap confidence intervals]), more than two times the estimated maximum sustainable yields, and almost three times what is estimated to be sustainable given current biomass levels. Therefore, annual catch estimates suggest that islanders are overfishing their coral reef fish stocks (Catch>Surplus). Estimated B_MMSY_ and the probability of being below BMMSY differed when using other surplus production models (Table S3). However, these did not impact the overall conclusion that fish stocks are expected to decline under the estimated level of overfishing.

**Figure 2.**
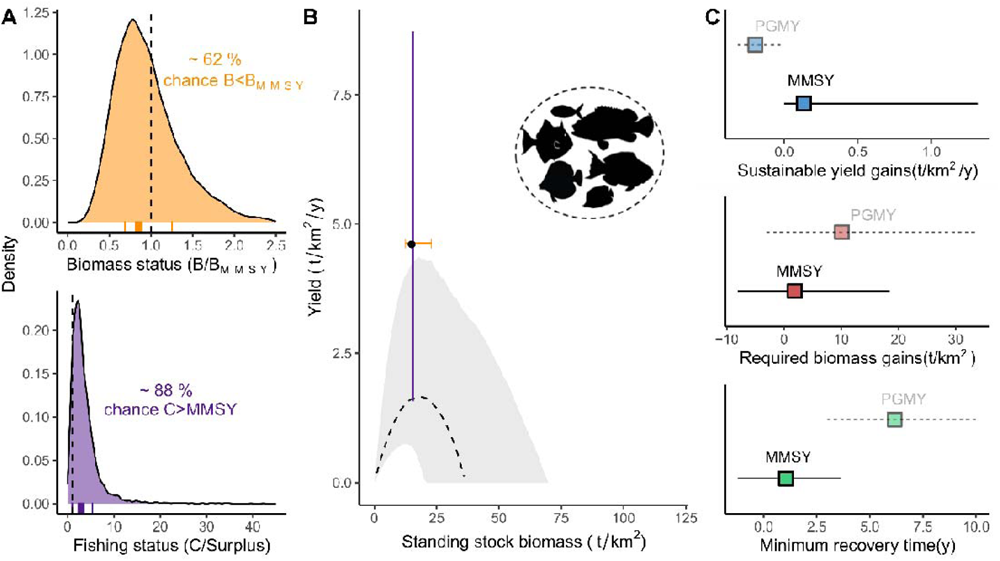
Whole assemblage sustainable reference points and assessment for the multispecies reef fishery of the study location using the Graham-Schaefer surplus production model. (A) Estimated multispecies maximum surplus production curve and assessment for our study location: Black line and grey ribbon represent the median and 90% uncertainty intervals for the estimated surplus production (t/km_2_/y) along a gradient of biomass. Overlaid point and error bars represent the per-unit-area median standing stock and annual catch estimates with their respective 95% adjusted bootstrap percentile intervals (orange for standing biomass and purple for annual catch). (B) Density distributions showing the probability of being below B_MMSY_ and of overfishing (accounting for uncertainty in the B_MMSY_ and surplus estimates from the global analysis applied to local conditions as well as the median biomass estimate for our study location). The dashed black vertical lines represent the limit between being “below B_MMSY_” and “not-below B_MMSY_” and the limit between being classified as “not-overfishing” and “overfishing”, respectively. Rug plots are the median status and 95% adjusted confidence intervals using the bootstrap samples (solid and dashed, respectively). (C) Potential yield gains, required biomass increases and minimum time required if reef fish stocks were recovered to B_MMSY_ or B_PGMY_ levels. Points are the medians and intervals are the 90% uncertainty intervals.

For the fishery to be sustainable at current biomass levels, catch would have to decrease around 3 t/km_2_/y, To maximize sustainable yields (i.e., reach B_MMSY_ from current biomass levels), reef fish biomass would need to increase by 2.0 [0– 18.6] t/km^2^ (median [90% uncertainty intervals]) and such recovery would take, in the most aggressive scenario (i.e., seascape moratorium), ∼1.1 [0– 6.8] years (Fig. 2C). This would allow sustainable catch to increase by ∼ 0.2 [0.1 – 1.3] t/km^2^/y, which is still substantially below current estimated catches, since current estimated annual catch is above the estimated MMSY. Lower gains (-0.2 [-0.6 – 0.7] t/km^2^/y), longer timeframes (6.2 [-0.5 – 17.4] y) and larger biomass increases (10.1 [-3.0 – 33.7] t/km^2^) would be required to reach more conservative targets, such as those that produce ‘pretty good multispecies yields‘ (B_PGMY,upper_; Fig. 2c).

Our status results were consistent with local fishers’ perceptions. We found that 83% of fishers perceived that the reef fish quantity and body length decreased over time, and that they spend more time looking for reef fish. Additionally, 67% of fishers reported that the species composition of their catch has changed, which they mostly (74%) attributed to reef fish getting too small or being difficult to find (Fig. 3).

**Figure 3.**
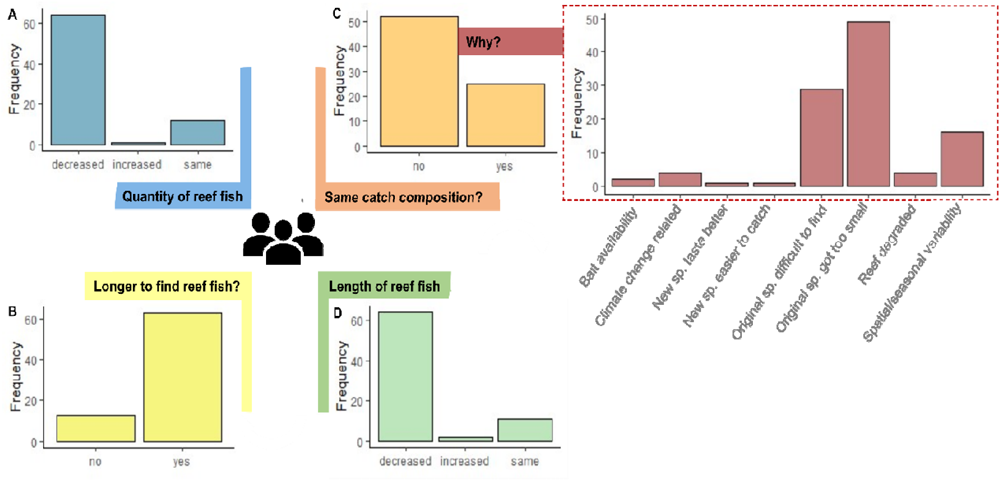
Fishers perceived reef fish stock status in comparison to the previous five years. **(A)** perceived change in the reef fish amount; (B) whether they had to spend more time to find and target reef fish; **(C)** whether the species composition of catch had changed and the reasons why fishers think the catch composition has changed; and (D) perceived change in the reef fish length.

## Discussion

Downscaling global models to local context can be a necessary step to sustainably manage and conserve natural resources (Cash and Moser 2000; McClanahan and Kosgei 2023), especially for locations with limited long-term monitoring. Here, we have constructed a framework to estimate locally relevant sustainable reference points and fishery assessments by integrating the results from global studies on coral reef fisheries when limited local fishery information is available. Our analyses revealed three key findings.

First, assessment results from our downscaled procedure were consistent with local fishers’ perceptions. Assessment results suggest that the reef fishery is unsustainable in comparison to whole assemblage proposed targets. Our analysis showed that median standing biomass values are currently lower than those needed to maximize yields, indicating that the reef fish assemblage is below levels that maximize production and fishers are overfishing their reef stocks. As overfishing takes place and a fishery becomes unsustainable, biomass is expected to become more depleted (Jackson et al. 2001), fish become more difficult to find and capture (McClanahan et al. 2008), and individual fish length becomes smaller on average (i.e., growth overfishing; Pauly 1994). Resource users, who are typically the first to notice and respond to these changes (e.g., by increasing effort or changing fishing locations; Silas et al. 2020), also reported these changes here, and their perceptions can thus help validate and triangulate global model outputs when integrated to local context (Neiss et al. 1999; Rochet et al. 2008; Ruano-Chamorro et al. 2017).

Second, local assessment results differ from those conducted at larger scales. Global assessments that use national reconstructed reef fish landings (Zeller et al. 2016) and reef area estimates (Spalding et al. 2001; UNEP-WCMC 2010) suggest overall that reefs in Papua New Guinea tend to be below B_MMSY_ but there is little overfishing (Zamborain-Mason et al. 2023). However, our results suggest that under current estimated local exploitation levels, the reef fish assemblage in our study location is expected to decline in the long-term. There are two main reasons for this discrepancy. One, spatial variability in environmental conditions, catch, or biomass within nations can impact what can be extracted sustainably, the level of exploitation, and the status of stocks, with average or median conditions (as provided by national-level analyses) not reflecting particular locations (e.g., Zeller et al. 2016; Karisa et al. 2020; McClanahan and Kosgei 2023). For example, in comparison to national estimates, Ahus had lower MMSY estimates due to higher-than-average SST and lower-than-average coral cover. Two, discordance between national and regional analyses could also reflect error or bias in jurisdiction-scale estimates such as representation of sampled reefs and timing, reef area, or catch estimates not being representative of ‘true’ local conditions. This finding emphasizes the importance of integrating local context and downscaling global models at appropriate scales for management.

Thirdly, we show that total annual catch estimates for our study location exceed not only what can be caught sustainably given the standing biomass (i.e., surplus), but also the estimated MMSY. This could mean one of three different things: (i) annual catch estimates for reef fish are not business-as-usual or biased high (e.g., sampled periods and fisher extrapolation are not representative of annual catch estimates); (ii) the estimated surplus production is biased low for this location (e.g., global model estimates are not representative of this region); or (iii) if our baseline assessment is accurate, then the population’s demand of reef fish is higher than the reef can provide, and reef fish populations are declining.

Accurate and precise annual catch estimates for most locations where reef fisheries operate are difficult to obtain (Russ 1991), especially if there are no monitoring programs in place (Teh et al. 2009). Here we obtained catch samples from different seasons and sexes and then extrapolated these with representative household surveys to get an annual catch estimate for the community. This approach did not discern among other factors such as age or experience, assuming the observed catch distribution for different sexes in different seasons is representative of their respective subpopulations. Changing this assumption if data are available (e.g., doing a weighted mean accounting experience, or fishing frequency), could be incorporated in future analyses that downscale global models to test additional catch estimates.

We used different surplus production models for the whole assemblage (Table S3). However, the estimated surplus production for the whole assemblage could be biased low for this location if either (i) the unfished biomass or/and the community biomass growth rate (e.g., Fig.1) were biased low (e.g., if the location was an outlier from global coral reefs in terms of how the assemblage responds to coral cover or recovery does not follow the same patterns as those inferred by space-for-time substitution, respectively) and/or (ii) the location’s surplus production does not follow the assumed dynamics (e.g., if species composition and species-specific interactions make the whole assemblage surplus production different to what global models infer). To overcome these in future work, it would be useful to apply global models to additional systems (e.g., that have recovery or validated reference point data), explore species-compositional changes and group-specific assessments and/or collect species-specific composition and additional temporal data in the study location (e.g., to evaluate whether reef assemblages decline as the assessment suggests).

A third option, consistent with fishers’ perceptions, is that our assessment results are representative of true conditions in which case stocks are expected to decline. While long-term monitoring will be needed to validate the expected decline in community biomass, the magnitude of the discrepancy between current estimated catch and sustainable reference points suggests that the community is unlikely to be able to meet its current fish demand even if stocks recover to maximum production levels. Thus, communities similar to our study location likely require alternative or diversified livelihoods and food supplies (e.g., mariculture, or increased access to offshore fisheries; Bell et al. 2018), and support mechanisms (e.g., financial help; Hilborn et al. 2005) to recover their reef fish stocks and return to sustainability. We estimated that stocks would take a median of ∼1.1 years to recover based on a complete seascape moratorium scenario, which would be devastating for communities that depend on reef resources for their daily food and income security. Indeed, in Ahus, completely restricting fishing would have significant impacts on the community and cause severe inequities, particularly for youth who have few skills or alternative livelihood options beyond fishing (Lau et al., 2021).

Most tropical reef fisheries around the globe remain unassessed. Typically, multispecies reef fisheries occur in the developing world, where research and formal management capacity are scarce (Worm and Branch 2012), and strong institutions to implement effective management measures are lacking (Hilborn et al. 2020). Many communities rely on co-management initiatives and inputs from non-governmental organizations (NGOs) to manage their fisheries, but clear reference points and monitoring to manage fisheries sustainably are often absent. While adequate assessment and effective management of reef resources will require long-term monitoring, validation, and adaptation (Free et al. 2019), we demonstrate how recent global models on reef fisheries can be downscaled to local context providing a pathway to perform baseline multispecies stock assessments for un-assessed fisheries around the globe at the scale of fisheries management. This will help provide baseline fishery assessments for un-assessed reef fisheries, and in turn, increase our understanding on how to update global models to make them more useful for resource practitioners and stakeholders.

## Acknowledgements

The authors would like to thank the Community of Ahus Island for their time, support, and knowledge; John Ben, John Kuange, Fidel Herry, Cristian Rojas, Sarah Sutcliffe and community members for field assistance and discussions; and Taryn Laubenstein for helping with image analyses. This research was supported by the Australian Research Council (CE140100020, FT160100047, FL230100201).

## Supporting Information

**Figure S1.**
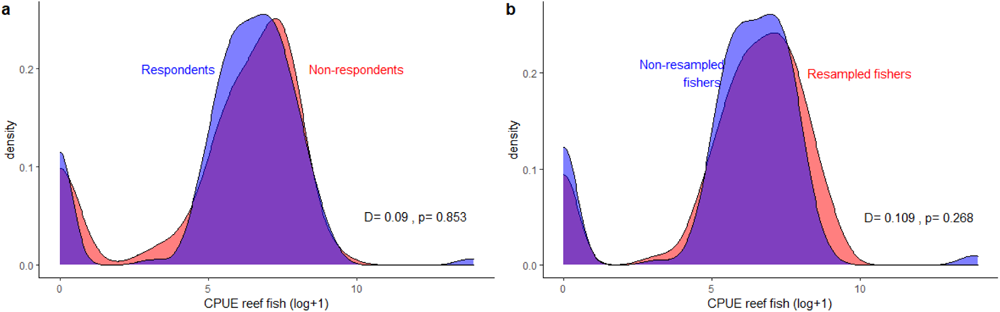
Comparison of catch per unit effort (CPUE) distributions from different sampled data. (a) those catch surveys whose fishers responded the long-term questions vs those that did not (i.e., non-respondents): and (b) catch surveys whose fishers were sampled more than once in the combined sapling period vs those fishers that were only sampled once. Black values at the bottom right are the Kolmogorov-Smirnov test results (i.e., D statistic and p-value). These show that there was no difference in the distributions of CPUE among groups (i.e., p-value>0.05).

**Figure S2.**
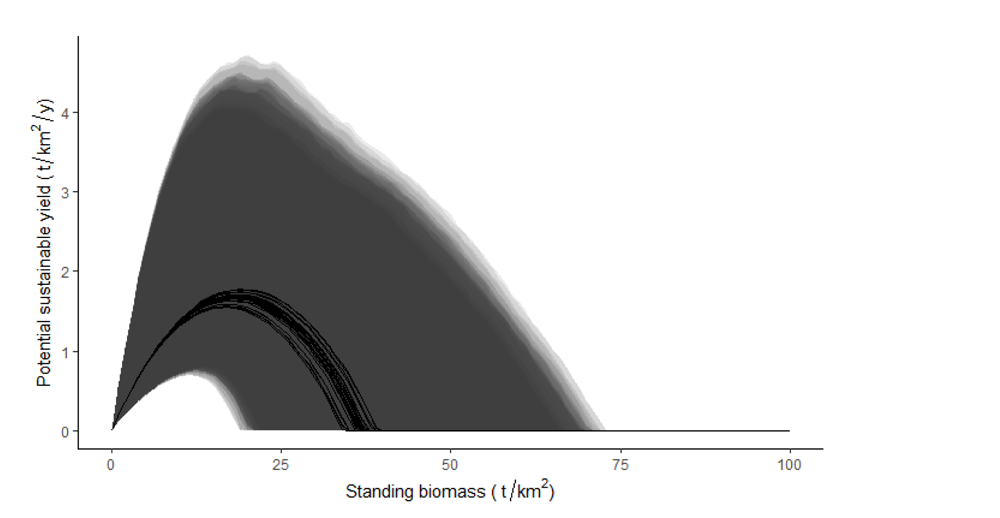
Surplus production curves for our study location for different sampled sites and years. Solid lines are the median estimated surplus along a gradient of biomass and polygons represent the 90% uncertainty intervals.

**Table S1.** Reef families included in our analyses.

| Fish family | Common family name |
| --- | --- |
| Acanthuridae | Surgeonfishes |
| Balistidae | Triggerfishes |
| Caesionidae | Fusiliers |
| Carangidae | Jacks |
| Chaetodontidae | Butterflyfishes |
| Cirrhitidae | Hawkfishes |
| Diodontidae | Porcupinefishes |
| Ephippidae | Batfishes |
| Haemulidae | Sweetlips |
| Kyphosidae | Drummers |
| Labridae | Wrasses and Parrotfishes |
| Lethrinidae | Emperors |
| Lutjanidae | Snappers |
| Monacanthidae | Filefishes |
| Mullidae | Goatfishes |
| Nemipteridae | Coral Breams |
| Pinguipedidae | Sandperches |
| Pomacanthidae | Angelfishes |
| Serranidae | Groupers |
| Siganidae | Rabbitfishes |
| Sparidae | Porgies |
| Sphyraenidae | Barracudas |
| Synodontidae | Lizardfishes |
| Tetraodontidae | Pufferfishes |
| Zanclidae | Moorish Idol |

**Table S2.**
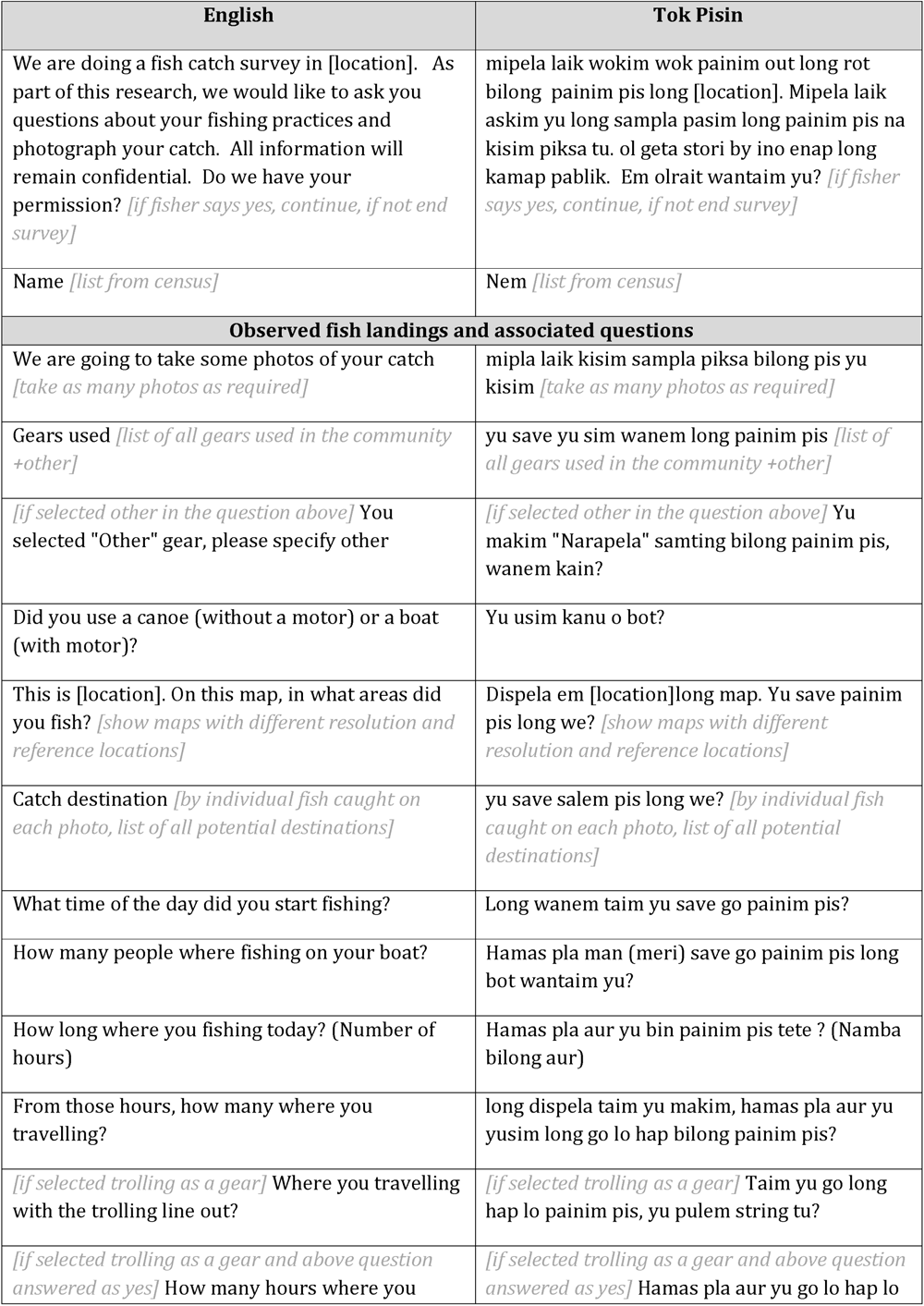
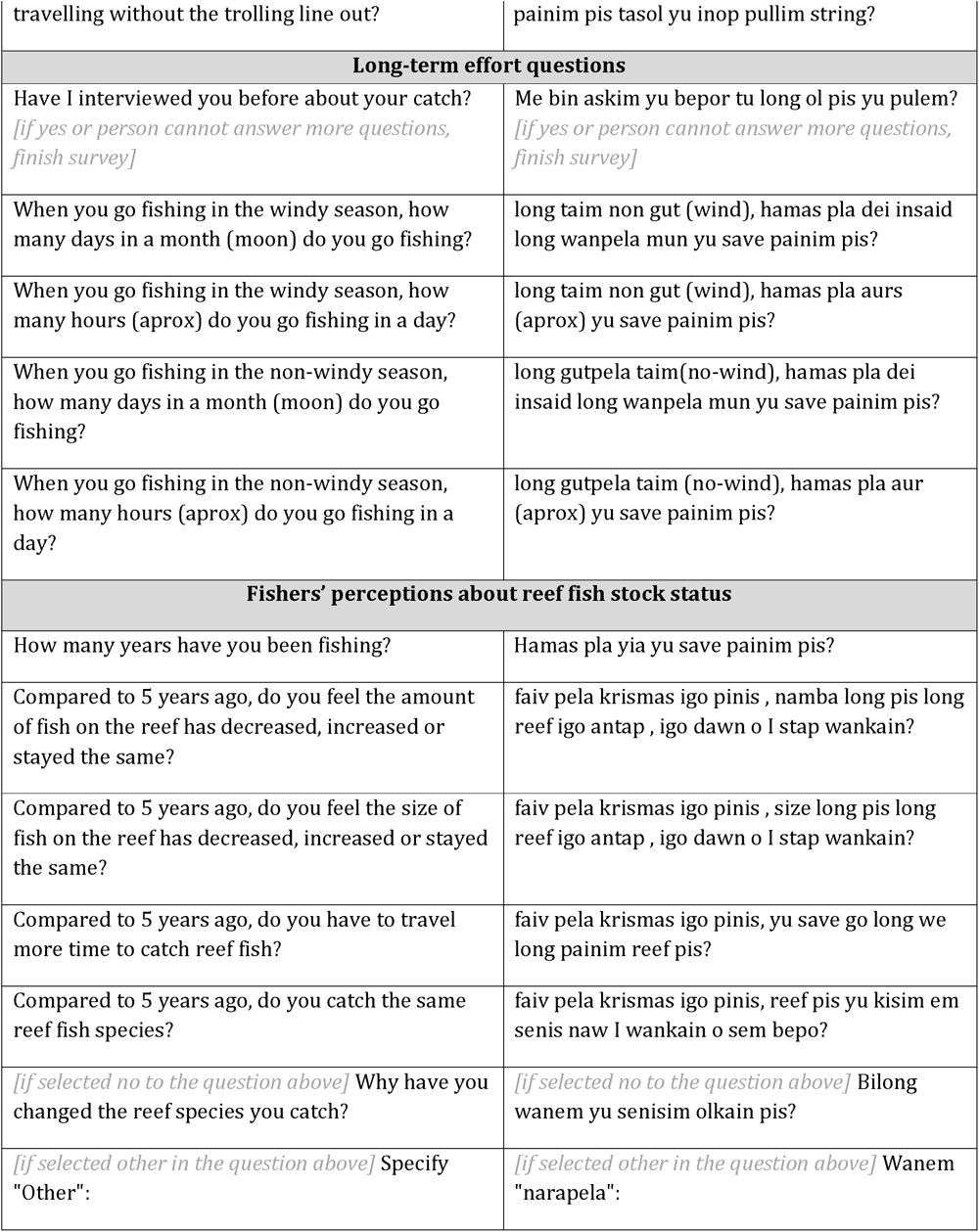
Catch survey questions.

| English | Tok Pisin |
| --- | --- |
| We are doing a fish catch survey in [location]. As part of this research, we would like to ask you questions about your fishing practices and photograph your catch. All information will remain confidential. Do we have your permission? *[if fisher says yes, continue, if not end survey]* | mipela laik wokim wok painim out long rot bilong painim pis long [location]. Mipela laik askim yu long sampla pasim long painim pis na kisim piksa tu. ol geta stori by ino enap long kamap pablik. Em olrait wantaim yu? *[if fisher says yes, continue, if not end survey]* |
| Name *[list from census]* | Nem *[list from census]* |
| Observed fish landings and associated questions |  |
| We are going to take some photos of your catch *[take as many photos as required]* | mipla laik kisim sampla piksa bilong pis yu kisim *[take as many photos as required]* |
| Gears used *[list of all gears used in the community +other]* | yu save yu sim wanem long painim pis *[list of all gears used in the community +other]* |
| *[if selected other in the question above]* You selected "Other" gear, please specify other | *[if selected other in the question above]* Yu makim "Narapela" samting bilong painim pis, wanem kain? |
| Did you use a canoe (without a motor) or a boat (with motor)? | Yu usim kanu o bot? |
| This is [location]. On this map, in what areas did you fish? *[show maps with different resolution and reference locations]* | Dispela em [location]long map. Yu save painim pis long we? *[show maps with different resolution and reference locations]* |
| Catch destination *[by individual fish caught on each photo, list of all potential destinations]* | yu save salem pis long we? *[by individual fish caught on each photo, list of all potential destinations]* |
| What time of the day did you start fishing? | Long wanem taim yu save go painim pis? |
| How many people where fishing on your boat? | Hamas pla man (meri) save go painim pis long bot wantaim yu? |
| How long where you fishing today? (Number of hours) | Hamas pla aur yu bin painim pis tete ? (Namba bilong aur) |
| From those hours, how many where you travelling? | long dispela taim yu makim, hamas pla aur yu yusim long go lo hap bilong painim pis? |
| *[if selected trolling as a gear]* Where you travelling with the trolling line out? | *[if selected trolling as a gear]* Taim yu go long hap lo painim pis, yu pulem string tu? |
| *[if selected trolling as a gear and above question answered as yes]* How many hours where you | *[if selected trolling as a gear and above question answered as yes]* Hamas pla aur yu go lo hap lo |
| travelling without the trolling line out? | painim pis tasol yu inop pullim string? |
| **Long-term effort questions** |  |
| Have I interviewed you before about your catch?<br>*[if yes or person cannot answer more questions, finish survey]* | Me bin askim yu bepor tu long ol pis yu pulem?<br>*[if yes or person cannot answer more questions, finish survey]* |
| When you go fishing in the windy season, how many days in a month (moon) do you go fishing? | long taim non gut (wind), hamas pla dei insaid long wanpela mun yu save painim pis? |
| When you go fishing in the windy season, how many hours (aprox) do you go fishing in a day? | long taim non gut (wind), hamas pla aurs (aprox) yu save painim pis? |
| When you go fishing in the non-windy season, how many days in a month (moon) do you go fishing? | long gutpela taim (no-wind), hamas pla dei insaid long wanpela mun yu save painim pis? |
| When you go fishing in the non-windy season, how many hours (aprox) do you go fishing in a day? | long gutpela taim (no-wind), hamas pla aur (aprox) yu save painim pis? |
| **Fishers' perceptions about reef fish stock status** |  |
| How many years have you been fishing? | Hamas pla yia yu save painim pis? |
| Compared to 5 years ago, do you feel the amount of fish on the reef has decreased, increased or stayed the same? | faiv pela krismas igo pinis, namba long pis long reef igo antap, igo dawn o I stap wankain? |
| Compared to 5 years ago, do you feel the size of fish on the reef has decreased, increased or stayed the same? | faiv pela krismas igo pinis, size long pis long reef igo antap, igo dawn o I stap wankain? |
| Compared to 5 years ago, do you have to travel more time to catch reef fish? | faiv pela krismas igo pinis, yu save go long we long painim reef pis? |
| Compared to 5 years ago, do you catch the same reef fish species? | faiv pela krismas igo pinis, reef pis yu kisim em senis naw I wankain o sem bepo? |
| *[if selected no to the question above]* Why have you changed the reef species you catch? | *[if selected no to the question above]* Bilong wanem yu senisim olkain pis? |
| *[if selected other in the question above]* Specify "Other": | *[if selected other in the question above]* Wanem "narapela": |

**Table S3.**
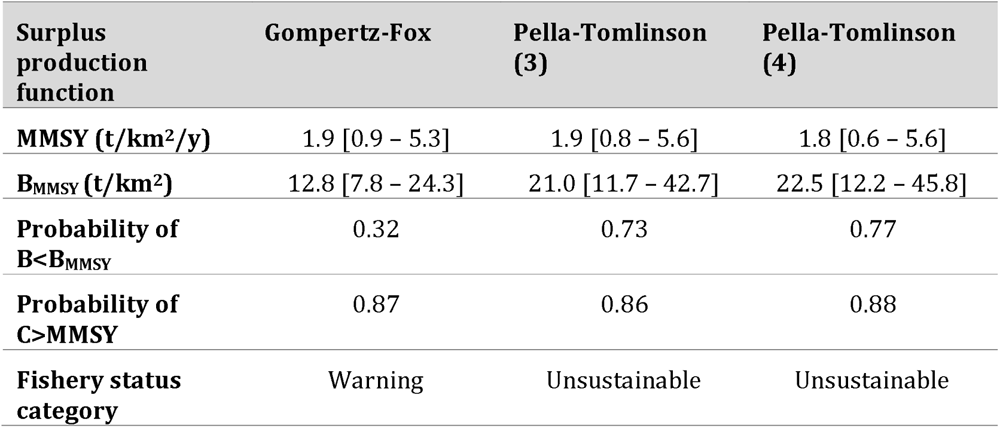
Estimated reference points and assessment results for our study location under alternate surplus production models. Reference points have medians and 90% uncertainty intervals (in brackets).

| Surplus<br>production<br>function | Gompertz-Fox | Pella-Tomlinson<br>(3) | Pella-Tomlinson<br>(4) |
| --- | --- | --- | --- |
| **MMSY (t/km<sup>2</sup>/y)** | 1.9 [0.9 – 5.3] | 1.9 [0.8 – 5.6] | 1.8 [0.6 – 5.6] |
| **B<sub>MMSY</sub> (t/km<sup>2</sup>)** | 12.8 [7.8 – 24.3] | 21.0 [11.7 – 42.7] | 22.5 [12.2 – 45.8] |
| **Probability of**<br>**B<B<sub>MMSY</sub>** | 0.32 | 0.73 | 0.77 |
| **Probability of**<br>**C>MMSY** | 0.87 | 0.86 | 0.88 |
| **Fishery status**<br>**category** | Warning | Unsustainable | Unsustainable |

